# Microevolutionary trends of tooth complexity in the Ibiza wall lizard, *Podarcis pityusensis*, mirror macroevolutionary trends in squamates

**DOI:** 10.1101/2025.05.14.653915

**Authors:** Stephanie Charlotte Woodgate, Alistair R. Evans, Bilal Kayed, Ana Pérez-Cembranos, Valentín Pérez-Mellado, Johannes Müller

## Abstract

Tooth complexity has long been known as a powerful indicator of feeding in mammals and reptiles, often used to infer diets of long-extinct species. However, how these trends are established below the species level remains unclear. Here, we use Orientation Patch Count Rotated (OPCR) to quantify tooth complexity across eight populations of *Podarcis pityusensis*, an insular omnivorous lizard endemic to the Pityusic Islands of the Western Mediterranean. These populations vary widely in their extent of herbivory and specialisation, which we use to assess the link between complexity and diet. We find that increasing herbivory is matched with increasing dental complexity, with comparable effect sizes to those found on the macroevolutionary scale. We do not, however, find clear evidence of divergence of tooth complexity between sexes within populations (in a way that would be consistent with niche divergence), meaning that the extent to which this trait is plastic remains controversial. Our results suggest that dental complexity is a highly powerful metric to infer diet, and an informative candidate for linking micro- and macroevolutionary patterns.

## BACKGROUND

A great outstanding controversy in evolutionary biology is overcoming the discrepancy often observed between evolutionary patterns on small taxonomic or time scales (microevolution) and those on large scales (macroevolution) (1–3). Linking macroevolutionary processes affecting phenotype with their microevolutionary underpinnings requires metrics that can be quantified at the population level, at the species level and beyond. More than simple identification of patterns, studying the specific functional implications of these metrics will elucidate how taxa interacting with their environment across scales shapes evolutionary trajectories. One such metric which has highly specific functional significance to the environment through association with diet is tooth shape. Herbivorous species tend to have more complex teeth than carnivorous species; this has been observed in diverse taxa, from mammals such as the order Carnivora, rodents and bats (4,5) to reptiles, where this relationship has been shown throughout squamates and even throughout non-avian reptiles (6–8). Similar patterns of complexity are found in toothless forms, such as the beaks of turtles (9) and grasshoppers (10). Indeed, this relationship is considered so strong that tooth shape has been used to infer diet of fossil reptiles and mammals (5,11–16). However, in their recent meta-analysis of tooth complexity according to diet, DeMers and Hunter conclude that the association between increasing herbivory and increasingly complex teeth cannot be generalised across amniotes, especially due to a lack of associations between diet and tooth complexity in some primates (17). Therefore, what may once have been considered received knowledge has recently been called into question.

Indeed, one aspect that has remained controversial is whether these findings can be translated onto the smaller, microevolutionary scale. Some species shift their dental shape through ontogeny; this is especially common in lizards, observed in diverse genera including *Amblyrhynchus, Ctenosaura, Varanus, Lacerta* and *Gallotia* (6,18–20), showing variation in tooth shape through development. However, less work has been done to ascertain to what extent differences in tooth shape among populations of the same species can be observed. In this regard, research has been mostly focussed on primate populations spanning large time or geographic scales, with studies varying on how much variation in tooth complexity is found and how strongly this can be linked to diet (21–23). In terms of studying contemporaneous populations, Vervust et al. found changes to the size of the teeth in a translocated population of *Podarcis siculus* lizards which consume more plants, without change in tooth shape (24). Naturally these works raise the question of how these broad-scale trends in tooth complexity are set up; specifically, whether tooth complexity (and other metrics such as tooth size and number) are able to shift on the population level to change with diet, or conserved at the species level.

Omnivorous species that consume different diets throughout their natural range are a great way to investigate these outstanding questions. Lacertid lizards living on islands are known for showing high degrees of omnivory (25); in particular, insular *Podarcis* lizards display highly variable diets, differing among populations of a single species, closely adapted to local resources (25–27). This makes them a brilliant candidate to investigate populational trends in tooth shape. In this work, we use *Podarcis pityusensis* as our candidate to investigate microgeographic trends in tooth complexity due to its diversity in phenotype and diet across its natural range of the Pityusic Islands in the Western Mediterranean. We choose this species because it is known to eat large proportions of plant and hard food, with some populations showing highly specialised diets (28).

We investigate complexity of teeth in a range of *P. pityusensis* populations with diverse diets to test the extent to which divergence in tooth morphology can be witnessed below the species level. We quantify tooth complexity using Orientation Patch Count (OPC) approaches, a powerful technique allowing fine-scale quantification of tooth topography which has been applied across the amniote tree of life (see (7,17,29) and references within). This work will operate under four hypotheses.

Our first hypothesis is that tooth complexity is underpinned by tooth size; particularly, that larger teeth have higher complexity. While it may be logical that in fabrication of the tooth, a larger surface area allows a greater level of complexity (see (30) and references within), this complexity-size relationship has been called into question on the developmental scale, with Harjunmaa et al. finding that cusp number is not necessarily increased by increasing tooth size (31).

Our second hypothesis is that diet drives the evolution of tooth complexity in *P. pityusensis*. Specifically, we hypothesise that there will be larger teeth and greater tooth complexity in populations that have a higher proportion of plants in the diet, as observed across reptiles (6,7) and many mammal species (4,5,32,33). Further, we hypothesise that populations that consume a greater percentage of ‘hard’ food (which require higher forces to crush) or plant food will have a higher tooth complexity, due to the need to break down material (4). We further hypothesise that there will be more variation in tooth complexity in populations with higher dietary diversity, as previously investigated by Selig et al., assessing to what extent the Niche Variation Hypothesis, that greater morphological complexity is observed in taxa with greater niche breadth (35), is applicable to teeth. We predict that the increase in complexity with plant material in the diet will not be linear; specifically, we predict that tooth complexity will increase more intensely at the very lowest levels of percentage herbivory; from animals consuming a very low proportion of plants (<20%) to plants becoming a moderate proportion of the diet (20-40%), as we propose tooth complexity to be the limiting factor in the amount of plant material that populations are able to exploit for food resources.

Under hypothesis three, we predict that diet will be the only factor controlling tooth complexity among populations of *P. pityusensis*, and that there will be no effect of ontogeny. Other lizard taxa display shifts in diet during ontogeny (18,19,36); however, this has so far not been reported for *P. pityusensis*.

Finally, to ascertain whether tooth shape is variable below the population level, we investigate whether complexity of teeth varies between sexes, as observed in a number of taxa (36). Dietary differences between sexes in omnivorous insular *Podarcis* species have previously been reported (37). Our fourth hypothesis predicts that divergence in diet between sexes drives divergence in tooth complexity between sexes. To test this prediction, we use a subset of populations from our dataset for which diet data was collected according to sex.

Overall, these hypotheses enable us to elucidate whether dietary differences drive divergence in tooth complexity in this lineage, allowing us to discern how variable tooth shape is in *P. pityusensis* lizards, and whether trends observed on the macroevolutionary level can also be demonstrated at the microevolutionary level.

## MATERIALS AND METHODS

### Specimen scanning and processing

The study species *Podarcis pityusensis* shows great diversity in diet throughout its natural range of the Balearic Islands. Eight populations with varying diets were selected for study, from each of which five specimens were included (Figure 1, full specimen list and information can be found in Supplementary Data Sheet 1, Dryad). Specimens were loaned from collections as following: specimens of Bleda Plana, Espartar, Es Vedrà and Trocadors populations from the University of Salamanca (USAL), specimens from Conillera from the Museum für Naturkunde Berlin (ZMB), specimens from Es Pouàs were loaned from Senckenberg Museum Frankfurt (SMF) and specimens from Espardell and Penjats were loaned from Museum Koenig Bonn (ZFMK). Specimens were identified as male or female either by the dimensions of the head or by femoral pore development and were also measured for Snout-Vent Length (SVL).

**Figure 1.**
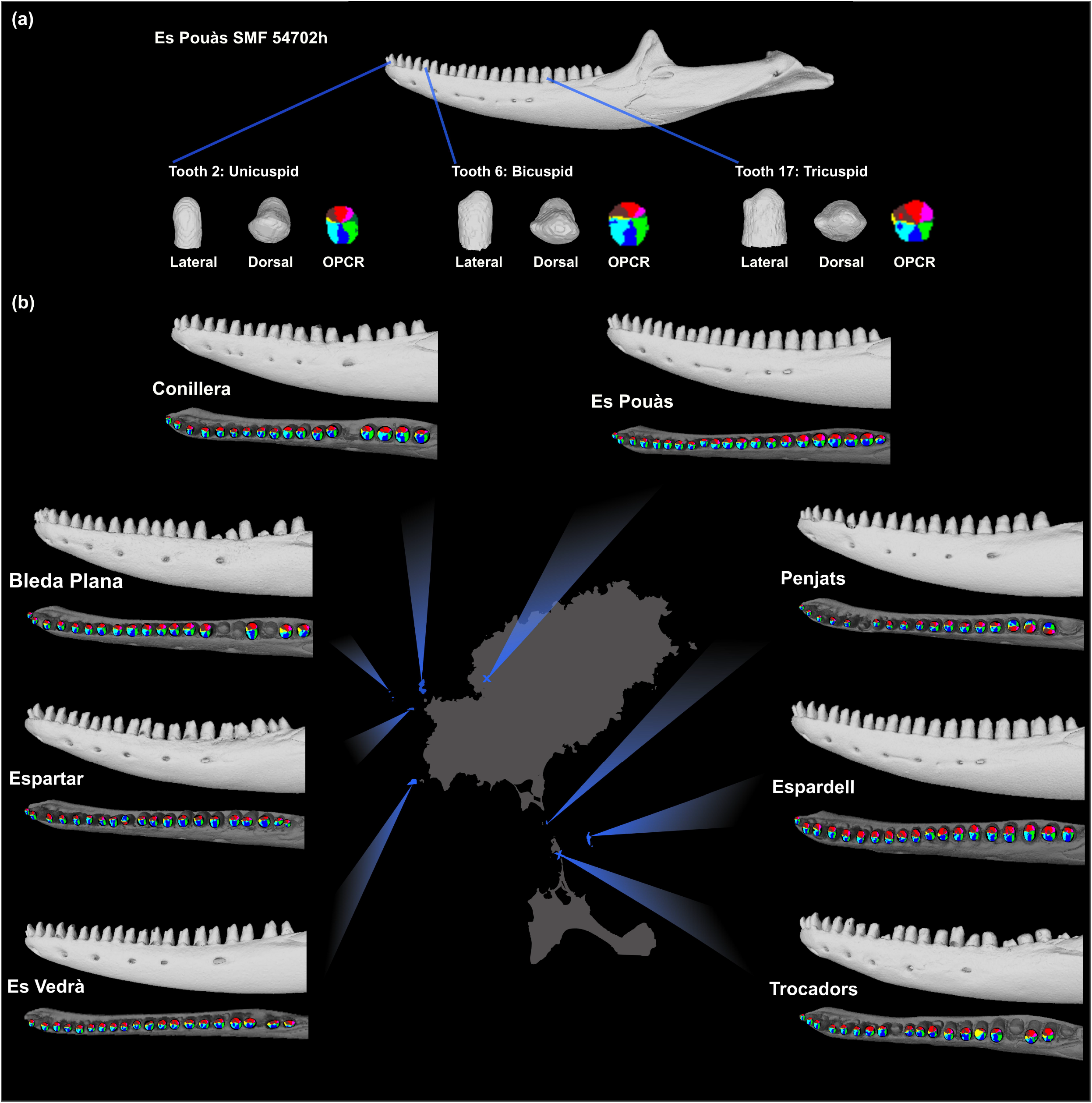
Tooth complexity by location. (a) Toothrow of specimen SMF 54702h, with shape and OPCR score of three example teeth. (b) Example toothrows from each population, with location specified in blue, showing jaws in lateral and dorsal aspect, not to scale. Colour of patches identified with OPCR analysis overlayed on dorsal view of the jaw.

Specimens were micro-CT scanned at the Museum für Naturkunde Berlin, then processed in VG Studio MAX; in some cases, this required segmenting upper and lower teeth. Left mandible models were then exported from VG Studios at the highest resolution available using the ‘grid-based watertight’ export function. High-resolution scans were processed in Meshlab to remove all mandible bone and unerupted or broken teeth, leaving a toothrow of left mandibular teeth to be investigated. The entire toothrow was investigated; however, in some cases teeth had to be removed from the analysis due to artificially flat surfaces caused by segmentation of tooth contacts; other teeth had smaller areas of cleaning with limited effect on the occlusal surface and were therefore included in the analysis, as specified in Supplementary Data Sheet 2. Individual teeth were oriented in Blender, with the occlusal surface facing in the positive z-axis and the anterior-most face of the tooth in the positive y-axis with the lingual (inner) surface of the tooth in the positive x-axis. An aligned mesh of each tooth was saved and co-ordinates defining the mesh of each tooth were then exported from Meshlab.

### Measuring tooth complexity

First, tooth complexity was measured by counting the number of cusps on each tooth. Teeth were counted according to six categories; monocuspid, bicuspid, tricuspid, rounded (in which no cusps can be identified, due to being ground down during use), broken and unerupted. The latter three categories were removed from further analysis.

Orientation Patch Count Rotation (OPCR) techniques were employed using the SurferManipulator (4) and Surfer 8 software, an approach based on Geographic Information System (GIS). OPCR is used here because it allows more sophisticated analysis of tooth complexity, identifying the number of faces on the surface of a tooth, rather than the cusp number alone, which means that more advanced metrics such as maximum tooth complexity and standard deviation in tooth complexity can be calculated. This quantifies tooth complexity by measuring the orientation (in cardinal and ordinal directions) of each co-ordinate in the surface grid. From this, the software identifies adjacent pixels on the tooth that have the same orientation; any group larger than two is defined as a ‘patch’; the software then gives an output of the number of patches, which is essentially the number of faces on the tooth surface (OPC). This is repeated at eight rotation intervals through a total of 45 degrees. The average of these eight orientations is then taken as the complexity value of that tooth (OPCR), to minimise the effect of orientation of the tooth on the patch number.

In order to standardise, each toothrow was re-gridded to 25 data rows per tooth, identified as the minimum number of rows for accurate quantification of tooth complexity of reptile teeth (7). This meant that some teeth too small for OPCR analysis (i.e., initially gridded with fewer than 25 data rows) were removed from the complexity analysis (Supplementary Data Sheet 2). In five specimens, five or more teeth were removed from the complexity analysis, either due to small size or flat surfaces from segmentation away from the upper teeth. Therefore, all analyses described below were re-run for a dataset excluding these five specimens; these results are included in Supplementary Data Tables 6-10.

Tooth size was measured in Meshlab by measuring the width of the base of the separated tooth perpendicular to the direction of the toothrow. This was done for all fully-erupted teeth, even those which had been removed from complexity analysis due to size. The entire length of the toothrow (from the first to the last tooth position, even if these teeth were missing or broken in the specimen) was also measured for each specimen, in order to be included in analyses to control for jaw size (described further below).

### Data processing

All data was imported into R for statistical analysis. The average number of cusps per tooth in the toothrow was calculated based only on teeth assessed as mono-, bi- or tri-cuspid. The OPCR data was used to calculate the mean complexity of each toothrow (mean complexity), the complexity of the most complex tooth (maximum complexity) and the standard deviation in tooth complexity (toothrow standard deviation).

The average cusp number per tooth should reflect the same information as the mean OPCR data, i.e both should reflect the average tooth complexity in that toothrow. To check this, an ANOVA was run testing the association between these two metrics.

The structure of the data was assessed. To check for normality, a Shapiro-Wilk test was performed, using the function shapiro.test() from the package *stats* (38). Next, to check for homogeneity of variances, a Levene’s test was performed, using the function leveneTest(). These were performed on the individual and average level for tooth complexity, tooth size and SVL.

### Diet data

Diet data (Supplementary Data Sheet 3) was gained from direct collection of scats in the field, analysed following the methods set out in (26). Percentage plant was taken directly while percentage ‘hard’ food was classified according the force needed to crush each item, with data taken from (27) and (40), and our own knowledge for any not included within (Supplementary Data Sheet 4). Hill q2 (the inverse Simpson index) was selected as the metric for dietary diversity. Dietary diversity was calculated using the function hill_taxa() from the package hillR (40). The significance of difference in dietary diversity among populations was analysed using the function mcpHill() from the package *simboot* (41).

### Data analysis

To investigate the relationship between tooth complexity and tooth size, a linear model linking OPCR to tooth length was generated, with toothrow length added as a covariate; this was done because raw tooth OPCR is not normally distributed. To check how size affects complexity on the level of the toothrow, ANCOVAs were performed (as all complexity data satisfied the assumptions of ANCOVA; see Supplementary Table 1), with mean tooth complexity as the dependent variable, mean tooth size as the independent variable and toothrow length as a covariate, using the function aov(). The same was repeated with maximum values. The relationship between tooth size and diet was investigated by setting up ANCOVAs of mean and maximum size against dietary metrics, with toothrow length as a covariate (see Supplementary Table 2).

To test the relationship between tooth complexity and diet, ANCOVAs were set up, with tooth complexity as the dependent variable, all dietary metrics as independent variables, alongside their interaction with tooth size and the covariate SVL included as a proxy for ontogeny. All specimens are adults, but it is possible that age may affect tooth shape, as witnessed in other lizard species (6,20). One model was set up for each tooth complexity metric: average number of cusps, mean tooth complexity, max tooth complexity, and toothrow standard deviation. A set of the same models with number of tooth positions in the toothrow as the proxy for ontogeny (42) were also set up (see Supplementary Table 3).

To assess the extent to which the observed increase in tooth complexity with percentage plant and hard material is continuous throughout the sample, we placed each dietary metric into categories, then tested the effect sizes of differences between each category. The amount of plant material in diets varies widely in our sample, from 10.0% (Bleda Plana) to 71.2% (Conillera). The percentage hard food in the diet varies less strongly in our dataset; from a low of 71.3% (Bleda Plana) to a high of 93.75% (Conillera). Four categories for degree of herbivory were assigned: very low herbivory defined as <20% plant material in the diet, low herbivory is defined as plant material 20-40%, medium herbivory is 40-60%, and high herbivory is >60% plant material. Cohen’s d, a measure of effect size that reflects how different one group is from another (43,44), was calculated using the function cohen.d() from the package *effsize* (45) between each contiguous group. Hedge’s g was then also computed, as a correction, often used for small sample sizes (46).

The same analysis was performed for percentage hard food, with five categories; very soft defined as <75% hard food, soft defined as hard food 75-80%, medium 80-85%, high 85-90%, and very high >90%.

Finally, within-population divergence in tooth complexity was investigated. All diet metrics were calculated for each sex at each location which had sufficient diet data by sex (i.e, more than one diet entry per sex); this included only three populations, Espardell, Es Vedrà and Penjats. Then, linear models were generated using the function lmer() from the package lmerTest (47) with location as a grouping factor, complexity metrics as the dependent variable and independent variables diet (in this case, specific diet for each sex) and sex, as well as the interaction between diet and sex as independent variables. ANOVAs were then run on these models using the function aov(). As this only includes three populations, it must be taken as a preliminary result.

Trends according to sex throughout the whole sample were also investigated, by performing Kruskal-Wallis tests using the function kruskal.test() to discern whether there were differences in tooth complexity and size between either sex. Diet data for each sex throughout the entire population was also calculated.

## RESULTS

Results of tests assessing the structure of the data can be found in Supplementary Table 1. As can be seen, number of clumps and raw tooth length violate the assumptions of ANOVA and ANCOVA.

Average cusp number is significantly associated with mean complexity from OPCR data (ANOVA F_(1, 38)_ = 11.02, adjusted R^2^ = 0.204, p = 0.002). This therefore shows that either method of measuring average complexity (whether counting the number of cusps or performing OPCR analysis) reflects similar information.

Complexity of the tooth is correlated with tooth size, when looking at the individual level; a linear model of OPCR by length with toothrow as a covariate is significant (F_(2, 633)_ = 3.51, adjusted R^2^ = 0.098, p = 2.41e^-15^, including tooth length with p = 1.05e^-15^ and toothrow length with p = 0.602). At the toothrow level, the average tooth size does not correlate with the average tooth complexity, whether measured via OPCR or cusp number, but the largest tooth size does correlate with the complexity of the most complex tooth (Table 1).

**Table 1.** Results from ANCOVAs investigating the relationship between tooth complexity and tooth size on the toothrow level. Asterisk indicates significant p-value.

| Model | Variable | Df | Sum Sq | Mean Sq | F value | Pr(>F) |
| --- | --- | --- | --- | --- | --- | --- |
| Average cusp number ~ mean tooth size + toothrow length | Mean size | 1 | 0.065 | 0.065 | 1.216 | 0.277 |
|  | Toothrow length | 1 | 0.114 | 0.114 | 2.133 | 0.153 |
|  | Residuals | 37 | 1.979 | 0.054 |  |  |
| Mean complexity ~ mean tooth size + toothrow length | Mean size | 1 | 0.003 | 0.003 | 0.007 | 0.936 |
|  | Toothrow length | 1 | 1.675 | 1.675 | 4.34 | 0.044* |
|  | Residuals | 37 | 14.293 | 0.386 |  |  |
| Maximum complexity ~ maximum tooth size + toothrow length | Max size | 1 | 6.340 | 6.340 | 7.272 | 0.011* |
|  | Toothrow length | 1 | 0.040 | 0.041 | 0.047 | 0.829 |
|  | Residuals | 37 | 32.260 | 0.872 |  |  |

Average tooth size and maximum tooth size increase with percentage plant material, but no other dietary metric (Supplementary Table 2). Average tooth complexity is significantly associated with percentage plant material in the diet, but not with any other dietary metric, which is true for mean cusp number and OPCR (Table 2). Maximum tooth complexity is not associated with any dietary metric, only with maximum tooth size. Standard deviation of tooth complexity shows no association with diet or size. These patterns remain when number of tooth positions are used as a proxy for ontogeny rather than SVL (Supplementary Table 3).

**Table 2.** Results from ANCOVAs investigating the tooth complexity, diet and tooth size, with SVL added as a covariate to control for ontogenetic stage. Asterisk indicates significant p-value.

|  | Variable | Df | Sum Sq | Mean Sq | F-value | P-value |
| --- | --- | --- | --- | --- | --- | --- |
| Average cusp number~ | Percent plant | 1 | 0.511 | 0.511 | 10.970 | 0.002* |
|  | Percent hard food | 1 | 0.027 | 0.027 | 0.582 | 0.451 |
|  | Dietary diversity | 1 | 0.001 | 0.001 | 0.021 | 0.885 |
|  | Max size | 1 | 0.003 | 0.003 | 0.071 | 0.791 |
|  | SVL | 1 | 0.017 | 0.017 | 0.373 | 0.546 |
|  | Percent plant:Max size | 1 | 0.010 | 0.010 | 0.211 | 0.649 |
|  | Percent hard food:Max size | 1 | 0.008 | 0.008 | 0.169 | 0.684 |
|  | Dietary diversity:Max size | 1 | 0.137 | 0.137 | 2.942 | 0.096 |
|  | Residuals | 31 | 1.444 | 0.047 |  |  |
| Mean tooth complexity~ | Percent plant | 1 | 2.393 | 2.393 | 6.119 | 0.019* |
|  | Percent hard food | 1 | 0.008 | 0.008 | 0.019 | 0.890 |
|  | Dietary diversity | 1 | 0.050 | 0.050 | 0.128 | 0.723 |
|  | Mean size | 1 | 0.185 | 0.185 | 0.473 | 0.497 |
|  | SVL | 1 | 0.085 | 0.085 | 0.217 | 0.644 |
|  | Percent plant:Mean size | 1 | 0.917 | 0.917 | 2.346 | 0.136 |
|  | Percent hard food:Mean size | 1 | 0.172 | 0.171 | 0.439 | 0.512 |
|  | Dietary diversity:Mean size | 1 | 0.038 | 0.038 | 0.098 | 0.756 |
|  | Residuals | 31 | 12.123 | 0.391 |  |  |
| Max tooth complexity~ | Percent plant | 1 | 1.642 | 1.642 | 1.770 | 0.193 |
|  | Percent hard food | 1 | 0.682 | 0.682 | 0.736 | 0.398 |
|  | Dietary diversity | 1 | 0.868 | 0.868 | 0.936 | 0.341 |
|  | Max size | 1 | 5.020 | 5.020 | 5.413 | 0.027* |
|  | SVL | 1 | 0.372 | 0.372 | 0.401 | 0.531 |
|  | Percent plant:Max size | 1 | 0.358 | 0.358 | 0.386 | 0.539 |
|  | Percent hard food:Max size | 1 | 0.802 | 0.802 | 0.865 | 0.360 |
|  | Dietary diversity:Max size | 1 | 0.147 | 0.147 | 0.159 | 0.693 |
|  | Residuals | 31 | 28.75 | 0.927 |  |  |
| Toothrow complexity standard deviation~ | Percent plant | 1 | 0.028 | 0.028 | 0.338 | 0.565 |
|  | Percent hard food | 1 | 0.070 | 0.070 | 0.853 | 0.363 |
|  | Dietary diversity | 1 | 0.030 | 0.030 | 0.369 | 0.548 |
|  | Mean size | 1 | 0.021 | 0.021 | 0.253 | 0.619 |
|  | SVL | 1 | 0.032 | 0.032 | 0.388 | 0.538 |
|  | Percent plant:Mean size | 1 | 0.000 | 0.000 | 0.005 | 0.942 |
|  | Percent hard food:Mean size | 1 | 0.027 | 0.027 | 0.324 | 0.573 |
|  | Dietary diversity:Mean size | 1 | 0.088 | 0.088 | 1.071 | 0.309 |
|  | Residuals | 31 | 2.534 | 0.082 |  |  |

Effect size analysis shows that the greatest difference is between groups of ‘low herbivory’ and ‘medium herbivory’; a ‘large’ difference is observed between these two groups, while a ‘very small’ difference is observed when transitioning between ‘very low’ to ‘low’ or ‘medium’ to ‘high’, classified according to (44) (Table 3). The effect size between ‘very low’ and ‘high’ herbivory is -0.78. Effect size analysis shows that the extent of difference between groups is the greatest at the lower end of hard food; a ‘large’ difference is seen between very soft and soft diet groups, a ‘large’ difference between soft and medium groups and a ‘medium’ difference between medium and hard, and hard and very hard groups (Supplementary Table 4).

**Table 3.** Showing results of tests investigating effect size between different categories of herbivore. Effect size classification is according to (44), based on Hedge’s g value.

| Extent of herbivory | Cohen's d | Hedges' g | Effect size classification |
| --- | --- | --- | --- |
| Very low:Low | 0.023 | 0.022 | Very small |
| Low:Medium | 0.873 | 0.836 | Large |
| Medium:High | 0.006 | 0.006 | Very small |
| Very low:High | -0.814 | -0.780 | Medium |

In the subsample employed to investigate drivers of tooth shape on the within-population level, dietary diversity and sex interact to shape the complexity of the most complex tooth (Table 4). This is despite the fact that maximum tooth complexity does not vary between sexes at the species level (KW-χ^2^= 0.451, p = 0.502) and also a general lack of divergence in dietary diversity between sexes, either within-population or the species level (Supplementary Table 5). Mean tooth complexity does not diverge between sexes either at the within-population level (see Table 4) or at the species level (cusp number: KW-χ^2^= 0.469, p = 0.493; OPCR data: KW-χ^2^= 0.0095, p = 0.922).

**Table 4.** Results of tooth complexity by diet and sex, using Type III Analysis of Variance Table with Satterthwaite’s method. Asterisk indicates significant p-value.

|  | Variable | Sum Sq | Mean Sq | NumDF | DenDF | F value | Pr(>F) |
| --- | --- | --- | --- | --- | --- | --- | --- |
| Average cusp number ~ | Percent plant | 0.003 | 0.003 | 1 | 4.090 | 0.082 | 0.788 |
|  | Percent hard food | 0.017 | 0.017 | 1 | 3.904 | 0.418 | 0.554 |
|  | Dietary diversity | 0.001 | 0.001 | 1 | 4.861 | 0.029 | 0.873 |
|  | Sex | 0.028 | 0.028 | 1 | 28.773 | 0.687 | 0.414 |
|  | Percent plant:Sex | 0.014 | 0.014 | 1 | 28.632 | 0.333 | 0.568 |
|  | Percent hard food:Sex | 0.015 | 0.015 | 1 | 28.679 | 0.370 | 0.548 |
|  | Dietary diversity:Sex | 0.018 | 0.018 | 1 | 28.585 | 0.433 | 0.516 |
| Mean tooth ~ | Percent plant | 0.122 | 0.122 | 1 | 4.054 | 0.350 | 0.586 |
|  | Percent hard food | 0.001 | 0.001 | 1 | 3.882 | 0.003 | 0.963 |
|  | Dietary diversity | 0.006 | 0.006 | 1 | 4.765 | 0.016 | 0.905 |
|  | Sex | 0.028 | 0.028 | 1 | 28.693 | 0.081 | 0.777 |
|  | Percent plant:Sex | 0.418 | 0.418 | 1 | 28.561 | 1.197 | 0.283 |
|  | Percent hard food:Sex | 0.075 | 0.075 | 1 | 28.605 | 0.215 | 0.646 |
|  | Dietary diversity:Sex | 0.003 | 0.003 | 1 | 28.517 | 0.010 | 0.922 |
| Max tooth ~ | Percent plant | 0.293 | 0.293 | 1 | 4.038 | 0.503 | 0.517 |
|  | Percent hard food | 0.023 | 0.023 | 1 | 3.977 | 0.039 | 0.853 |
|  | Dietary diversity | 0.473 | 0.473 | 1 | 4.279 | 0.812 | 0.415 |
|  | Sex | 0.446 | 0.446 | 1 | 28.250 | 0.767 | 0.389 |
|  | Percent plant:Sex | 0.001 | 0.001 | 1 | 28.206 | 0.002 | 0.962 |
|  | Percent hard food:Sex | 0.093 | 0.093 | 1 | 28.220 | 0.159 | 0.693 |
|  | Dietary diversity:Sex | 2.624 | 2.624 | 1 | 28.192 | 4.504 | 0.043* |

Analyses excluding specimens that had five or more teeth excluded from OPCR analysis returned similar results with similar p-values as described above; a linear model of mean tooth complexity to average cusp number returned F_(1,33)_= 12.23, Adjusted R-squared: 0.2483, p= 0.00137, other results reported in Supplementary Tables 6-10. However, the analysis of effect size was different, giving different effect size classifications; particularly, a ‘large’ effect size of -0.95 for the difference between lowest and highest herbivory (Supplementary Tables 9-10).

## DISCUSSION

Our results indicate that size of the largest tooth in the toothrow is correlated with its complexity; the larger the largest tooth, the more complex the most complex tooth. This trend does not hold for average size and average complexity, suggesting that the surface area is only important in determining OPCR once high complexity is reached. This is a trend which may be difficult to assess while quantifying complexity at the level of cusp number (31), and shows the importance of using techniques which can quantify complexity on a finer level, such as OPCR. Increasing tooth size driving increasing complexity is observed in multituberculates (16) and other lizard species such as *Amblyrhynchus cristatus* (6); however, these changes relate to the level of averages rather than most complex or largest. Therefore, we can accept hypothesis one, in *P. pityusensis*, tooth complexity is underpinned by tooth size, but only via the largest tooth sizes allowing the highest complexities to be achieved.

Increasing amount of plant food in the diet is correlated with increasing complexity and size in the teeth of *P. pityusensis*, allowing us to accept hypothesis two. This is in contrast to trends in *Podarcis siculus* in which lizards from a more herbivorous islet population have larger average tooth size with difference in complexity (measured via number of cusps) (24). Instead, our results fit more clearly with trends of increasing complexity with increasing herbivory observed on the macroevolutionary scale, across squamates and other non-avian reptiles (6,7). This relationship is not affected by ontogeny according to either of the ontogeny metrics we investigated, allowing us to accept hypothesis three. We do not find a relationship between the percentage hard food in the diet and either maximum or mean tooth complexity in the sample. These findings, alongside the fact that hard food does not drive bite force evolution in *P. pityusensis* populations (*submitted*) therefore suggest that diets which are generally ‘hard’ or ‘soft’ do not impose strong selective pressures in *P. pityusensis*; it is specifically the proportion of plant food in the diet which drives phenotypic evolution.

We further find a lack of correlation between disparity in tooth complexity and dietary diversity, similar to findings in primates (34). This therefore suggests that the selective pressures imposed on tooth morphology by the diet mostly focus on the ability to process the food material. More diverse teeth are not needed to consume more diverse diets; diet is imposing selective pressures only according to the need to process the amount of plant food. This is the case even despite limited food processing in the *P. pityusensis* jaw compared to other taxa such as mammals.

Effect size analysis suggests that the most significant changes in dental complexity occur between two groups of ‘middle extent’ of herbivory, from <40% herbivory to 40%-60% herbivory. The changes here may reflect an ecological shift, e.g from relying on plants as a supplementary part of the diet to incorporating plants as a significant part of daily food consumption. The ‘supplementary’ diet imposes weak or negligible selective pressures on tooth complexity while the regular consumption of plants imposes stronger selective constraints. Indeed, the negligible increase in tooth complexity from <60% plant material to >60% suggests that strong selective pressures are already in place at the medium level of herbivory.

The ‘fallback food’ phenomenon could be an explanation for this. Populations of *Podarcis lilfordi* (sister species to *Podarcis pityusensis*) living on Aire Island rely on *Helicodiceros muscivorus* fruits for food during times of low animal prey availability, especially years with very low rainfall (48). *P. lilfordi* thus has a ‘fallback’ diet of fruits, which supplements the regular omnivorous diet during unfavourable environmental conditions. Our finding that *P. pityusensis* populations with very low and low herbivory have similar tooth complexity suggests that all populations have teeth complex enough to process some plant material, perhaps because *P. pityusensis* may also need to ‘fallback’ on plant diet during tough environmental conditions. Combined with findings in *P. lilfordi*, this may suggest that all Balearic *Podarcis* populations are adapted to include plant material as a supplementary part of the diet.

DeMers and Hunter found that in groups where herbivores have more complex teeth than faunivores, the extent of increase in complexity in association with plant diet varies from taxon to taxon (17). The authors find that the increase in complexity from insectivore to herbivore gives a Hedge’s g value of 1.05 across squamates (data from (6)) and 0.91 across all toothed saurians (data from (7)), while our study reports a comparable, but slightly lower, value of 0.78 (see Table 1, minimal:high); however, in the reduced dataset of 35 specimens, we found an SMD of -0.95 between minimal to high herbivory (see Supplementary Table 9). These highly comparable values of effect size are especially interesting because they are despite the fact that true herbivory (defined as >90% plant food, which is witnessed in only 0.8% of lizard species (49)) is not observed in any *P. pityusensis* populations; indeed, no *P. pityusensis* populations display post-cranial adaptations to herbivory (28). We therefore support the assertion of DeMers and Hunter that OPCR data can be used to infer diet in reptiles (17); indeed, the fact that microevolutionary trends in tooth complexity follow the same patterns as macroevolutionary trends suggests that studying tooth complexity using OPCR data is a powerful way to infer diet in reptiles. We would even go as far as suggesting that this approach can be used across amniotes, with the possible exception of some primates.

Teeth have previously been found to be a useful link between micro- and macroevolution in primates (50), and our data suggests this is also true in squamates. A central question posed by our findings is to what extent the differences in tooth complexity are phenotypically variable or evolutionarily fixed. While dietary diversity does interact with sex to show higher complexity of the most complex tooth at the within-population level, dietary diversity does not drive complexity of the most complex tooth at the among-population level. We find a lack of relationship of tooth complexity, sex and herbivory in *P. pityusensis* despite percentage plant material in the diet being higher in male lizards, leading us to reject hypothesis four. This may suggest that evolutionary patterns identified at the among-population level cannot be linked to those observed at the within-population level; however, our within-population analysis has a very limited sample size, so must be considered a preliminary result.

Investigating within-population trends further could be a useful future avenue of this work.

Each population tested here originates from a different island or islet of the Pityusic Islands, and their genetic interrelatedness is as-yet controversial (51,52). Harjunmaa et al. found evidence for what they term a ‘general bias against increase in dental complexity’, suggesting that increases in complexity are slow and rely on multiple alterations to developmental factors (31). However, the picture may be more complex in reptiles: Lafuma et al. find evidence for at least 24 transitions towards multi-cuspidness across the squamate tree (8). Particularly, these authors hypothesise a ‘correlated progression’ model between tooth complexity and plant consumption, which may have facilitated diversification in clades such as *Podarcis*. Our results suggest this process may even be taking place on the microevolutionary scale in *Podarcis pityusensis*; however, the extent to which the trends we observe are genetically based or developmentally plastic require further study. In any case, this work shows the clear efficacy in using tooth complexity metrics to investigate diet, both below the species level and at the macroevolutionary level. We therefore add further evidence that tooth complexity is a powerful metric through which to link microevolutionary and macroevolutionary patterns from the population level to the order level, and perhaps even beyond.

## CONCLUSION

We show that microevolutionary trends of tooth complexity of *Podarcis pityusensis* consuming different diets are the same as those observed among saurians on the macroevolutionary scale, with increasing tooth complexity associated with increasing amount of plant food in the diet. Indeed, the effect size between high and lowest herbivory is comparable within *P. pityusensis* and across non-avian reptiles, despite the fact that there are no ‘true’ herbivores within our sample. The greatest change in tooth complexity occurs between populations which have a ‘low’ and ‘medium’ extent of plant consumption, suggesting that the strongest selective influence of herbivory on tooth shape is when plants become a regular part of the diet, and possibly suggesting that all populations rely on plants as a ‘fallback’ food during periods of lower prey availability. We find no evidence of average tooth shape varying below the population level (i.e, between the sexes); however, those results must be considered preliminary due to the limited size of our dataset. We conclude that tooth complexity is a highly powerful metric to infer diet in reptiles, and perhaps amniotes as a whole; and further, a highly powerful metric to link evolutionary trends over micro- and macro-scales.

## Supporting information

Supplementary Results Tables

## ACKNOWLEDGEMENTS

We would like to thank Linda Mogk from the Senckenberg Museum Frankfurt (SMF), Claudia Koch and Morris Flecks from Museum Koenig Bonn (ZFMK) for loaning specimens included in this study. We would like to thank Kristin Mahlow-Tillack (MfN) for micro-CT scanning specimens, and Frank Tillack (MfN) for providing access to specimens and for help in preparation of specimens for scanning. Thanks to Vincent Fantino (MfN) for help preparing 3D models from micro-CT scans.

