## Supplementary Results Tables for "Microevolutionary trends of tooth complexity in the Ibiza wall lizard, *Podarcis pityusensis*, mirror macroevolutionary trends in squamates"

**Supplementary Data to accompany the manuscript 'Microevolutionary trends of tooth complexity in the Ibiza wall lizard, *Podarcis pityusensis,* mirror macroevolutionary trends in squamates,**

| | Shapiro-Wilk test | | Levene Test | |
|---|---|---|---|---|
| **Variable** | **W** | **P-value** | **F value** | **P-value** |
| Average cusp number | 0.9892 | 0.9633 | 0.2962 | 0.9503 |
| Clump number | 0.97002 | 3.854e-10* | 2.4962 | 0.01554* |
| Raw tooth length | 0.9671 | 1.305e-11* | 2.0111 | 0.05138 |
| Mean complexity | 0.98268 | 0.7875 | 0.7095 | 0.6642 |
| Maximum complexity | 0.95025 | 0.07733 | 0.6808 | 0.687 |
| Mean tooth size | 0.98975 | 0.9713 | 0.4629 | 0.8538 |
| Max tooth size | 0.99126 | 0.9873 | 0.3638 | 0.9165 |
| SVL | 0.98492 | 0.862 | 0.7129 | 0.6615 |

Supplementary Table 1. Results of Shapiro-Wilk tests for normality and Levene Tests for homogeneity. Asterisk indicates significant p-value.

| | Variable | Df | Sum Sq | Mean Sq | F value | Pr(>F) |
|---|---|---|---|---|---|---|
| Mean size ~ | Percent plant | 1 | 0.00305 | 0.00305 | 8.561 | 0.006* |
| | Percent hard food | 1 | 0.00016 | 0.00016 | 0.44 | 0.512 |
| | Dietary diversity | 1 | 0.00002 | 0.00002 | 0.052 | 0.82 |
| | Toothrow length | 1 | 0.03849 | 0.03849 | 108.052 | 3.05E-12* |
| | Residuals | 35 | 0.01247 | 0.00036 | | |
| Max size ~ | Percent plant | 1 | 0.01514 | 0.01514 | 12.576 | 0.00113* |
| | Percent hard food | 1 | 0.00138 | 0.00138 | 1.143 | 0.29236 |
| | Dietary diversity | 1 | 0.0009 | 0.0009 | 0.744 | 0.39421 |
| | Toothrow length | 1 | 0.06264 | 0.06264 | 52.024 | 2.03E-08* |
| | Residuals | 35 | 0.04214 | 0.0012 | | |

Supplementray Table 2. Results from ANCOVAs of tooth size against diet, with variables specified. Asterisk indicates significant p-value.

| | Variable | Df | Sum Sq | Mean Sq | F-value | P-value |
|---|---|---|---|---|---|---|
| Average cusp number ~ | Percent plant | 1 | 0.511 | 0.511 | 10.794 | 0.00253* |
| | Percent hard food | 1 | 0.0271 | 0.0271 | 0.573 | 0.45491 |
| | Dietary diversity | 1 | 0.001 | 0.001 | 0.021 | 0.88632 |
| | Mean size | 1 | 0.0061 | 0.0061 | 0.13 | 0.72099 |
| | Tooth positions | 1 | 0.0014 | 0.0014 | 0.029 | 0.86588 |
| | Percent plant: Mean size | 1 | 0.0382 | 0.0382 | 0.807 | 0.37585 |
| | Percent hard food: Mean size | 1 | 0.003 | 0.003 | 0.063 | 0.80367 |
| | Dietary diversity: Mean size | 1 | 0.103 | 0.103 | 2.176 | 0.15027 |
| | Residuals | 31 | 1.4677 | 0.0473 | | |
| Mean tooth complexity~ | Percent plant | 1 | 2.393 | 2.3928 | 6.165 | 0.0187* |
| | Percent hard food | 1 | 0.008 | 0.0075 | 0.019 | 0.89 |
| | Dietary diversity | 1 | 0.05 | 0.0501 | 0.129 | 0.7217 |
| | Mean size | 1 | 0.185 | 0.185 | 0.477 | 0.4951 |
| | Tooth positions | 1 | 0.205 | 0.2055 | 0.529 | 0.4723 |
| | Percent plant: Mean size | 1 | 0.841 | 0.8412 | 2.167 | 0.1511 |
| | Percent hard food: Mean size | 1 | 0.211 | 0.2114 | 0.545 | 0.466 |
| | Dietary diversity: Mean size | 1 | 0.044 | 0.0444 | 0.114 | 0.7375 |
| | Residuals | 31 | 12.033 | 0.3881 | | |
| Max tooth complexity~ | Percent plant | 1 | 1.642 | 1.642 | 1.782 | 0.1916 |
| | Percent hard food | 1 | 0.682 | 0.682 | 0.741 | 0.3961 |

| | | | | | | |
|---|---|---|---|---|---|---|
| | Dietary diversity | 1 | 0.868 | 0.868 | 0.943 | 0.3391 |
| | Max size | 1 | 5.02 | 5.02 | 5.45 | 0.0262* |
| | Tooth positions | 1 | 1.104 | 1.104 | 1.199 | 0.282 |
| | Percent plant: Max size | 1 | 0.083 | 0.083 | 0.09 | 0.7659 |
| | Percent hard food: Max size | 1 | 0.57 | 0.57 | 0.619 | 0.4375 |
| | Dietary diversity: Max size | 1 | 0.115 | 0.115 | 0.125 | 0.7262 |
| | Residuals | 31 | 28.557 | 0.921 | | |
| Toothrow standard deviation~ | Percent plant | 1 | 0.0277 | 0.02766 | 0.362 | 0.552 |
| | Percent hard food | 1 | 0.0697 | 0.06973 | 0.911 | 0.347 |
| | Dietary diversity | 1 | 0.0302 | 0.03019 | 0.394 | 0.535 |
| | Mean size | 1 | 0.0207 | 0.02067 | 0.27 | 0.607 |
| | Tooth positions | 1 | 0.198 | 0.19799 | 2.587 | 0.118 |
| | Percent plant: Mean size | 1 | 0.031 | 0.03102 | 0.405 | 0.529 |
| | Percent hard food: Mean size | 1 | 0.0056 | 0.00556 | 0.073 | 0.789 |
| | Dietary diversity: Mean size | 1 | 0.0733 | 0.07331 | 0.958 | 0.335 |
| | Residuals | 31 | 2.3722 | 0.07652 | | |

Supplementary Table 3. Results from ANCOVAs investigating the tooth complexity, diet and tooth size, with tooth positions added as a covariate to control for ontogenetic stage. Asterisk indicates significant p-value.

| Amount of hard food | Cohen's d | Hedges' g | Effect size classification |
|---|---|---|---|
| Very soft: Soft | 1.409832 | 1.3269 | Large |
| Soft: Medium | 1.007795 | 0.9485127 | Large |
| Medium: High | 0.8232009 | 0.7884178 | Medium |
| High: Very high | 0.554668 | 0.5312313 | Medium |
| Very soft: Very high | 1.424282 | 1.286448 | Large |

Supplementary Table 4. Showing results of tests investigating effect size between different categories of hardness of diet. Effect size classification is according to Sawilowsky, (2009), based on Hedge's g value.

| Population | Dietary metric | Male | Female | P-value difference between sexes |
|---|---|---|---|---|
| TOTAL | Percentage plant | 44.96296296 | 14.72222222 | |
| | Percentage hard food | 83.05084746 | 72.97297297 | |
| | Inverse Simpson Index | 4.574244 | 6.194570 | 0.2460 |
| Espardell | Percentage plant | 93.33333333 | 15 | |
| | Percentage hard food | 71.42857143 | 75 | |
| | Inverse Simpson Index | 3.769231 | 4.500000 | 0.9004 |
| Es Vedrà | Percentage plant | 0 | 2 | |
| | Percentage hard food | 78.94736842 | 77.77777778 | |
| | Inverse Simpson Index | 4.945205 | 5.400000 | 0.8978 |
| Penjats | Percentage plant | 0 | 0 | |
| | Percentage hard food | 100 | 70 | |
| | Inverse Simpson Index | 3.571429 | 7.142857 | 1.0000 |

Supplementary Table 5. Results from within-population (i.e, between sex) analysis dietary divergence. P-values are the results of the mcpHill() test based on the Inverse Simpson index between the two groups, with Tukey type matrix which takes into account multiple entries.

| Diet metric | Variable | Df | Sum Sq | Mean Sq | F value | Pr(>F) |
|---|---|---|---|---|---|---|
| Average cusp number ~ mean tooth size + toothrow length | Mean size | 1 | 0.0779 | 0.07795 | 1.436 | 0.24 |
| | Toothrow length | 1 | 0.1248 | 0.1248 | 2.298 | 0.139 |

| | | | | | | |
|---|---|---|---|---|---|---|
| | Residuals | 32 | 1.7375 | 0.0543 | | |
| Mean complexity ~ mean tooth size + toothrow length | Mean size | 1 | 0.048 | 0.0482 | 0.129 | 0.722 |
| | Toothrow length | 1 | 1.36 | 1.3597 | 3.646 | 0.0652 |
| | Residuals | 32 | 11.934 | 0.3729 | | |
| Maximum complexity ~ maximum tooth size + toothrow length | Max size | 1 | 6.136 | 6.1362 | 6.454 | 0.0161* |
| | Toothrow length | 1 | 0.058 | 0.058 | 0.061 | 0.8067 |
| | Residuals | 32 | 30.424 | 0.951 | | |

Table 6. Results from a reduced dataset of 35 toothrows: ANCOVAs investigating the relationship between tooth complexity and tooth size on the toothrow level. Asterisk indicates significant p-value.

| | Variable | Df | Sum Sq | Mean Sq | F-value | P-value |
|---|---|---|---|---|---|---|
| Average cusp number~ | Percent plant | 1 | 0.4811 | 0.4811 | 11.459 | 0.00227* |
| | Percent hard food | 1 | 0.0771 | 0.0771 | 1.836 | 0.18705 |
| | Dietary diversity | 1 | 0.0105 | 0.0105 | 0.249 | 0.62187 |
| | Max size | 1 | 0.0093 | 0.0093 | 0.222 | 0.64153 |
| | SVL | 1 | 0.0471 | 0.0471 | 1.122 | 0.29933 |
| | Percent plant: Max size | 1 | 0.0017 | 0.0017 | 0.04 | 0.84278 |
| | Percent hard food: Max size | 1 | 0.106 | 0.106 | 2.525 | 0.12413 |
| | Dietary diversity: Max size | 1 | 0.116 | 0.116 | 2.764 | 0.1084 |
| | Residuals | 26 | 1.0915 | 0.042 | | |
| Mean tooth complexity~ | Percent plant | 1 | 2.642 | 2.6418 | 8.034 | 0.00876* |
| | Percent hard food | 1 | 0.042 | 0.0422 | 0.128 | 0.72307 |
| | Dietary diversity | 1 | 0.012 | 0.0119 | 0.036 | 0.85064 |
| | Max size | 1 | 0.024 | 0.0239 | 0.073 | 0.78978 |
| | SVL | 1 | 0.013 | 0.0128 | 0.039 | 0.84524 |
| | Percent plant: Max size | 1 | 1.321 | 1.3215 | 4.019 | 0.05551 |
| | Percent hard food: Max size | 1 | 0.701 | 0.701 | 2.132 | 0.15625 |
| | Dietary diversity: Max size | 1 | 0.037 | 0.0374 | 0.114 | 0.73853 |
| | Residuals | 26 | 8.549 | 0.3288 | | |
| Maximum tooth complexity~ | Percent plant | 1 | 1.381 | 1.381 | 1.343 | 0.257 |
| | Percent hard food | 1 | 0.471 | 0.471 | 0.458 | 0.5046 |
| | Dietary diversity | 1 | 1.248 | 1.248 | 1.213 | 0.2808 |
| | Max size | 1 | 4.778 | 4.778 | 4.646 | 0.0406* |
| | SVL | 1 | 0.254 | 0.254 | 0.247 | 0.6231 |
| | Percent plant: Max size | 1 | 0.114 | 0.114 | 0.111 | 0.7417 |
| | Percent hard food: Max size | 1 | 1.626 | 1.626 | 1.581 | 0.2198 |
| | Dietary diversity: Max size | 1 | 0.009 | 0.009 | 0.008 | 0.928 |
| | Residuals | 26 | 26.737 | 1.028 | | |
| Toothrow standard deviation~ | Percent plant | 1 | 0.0143 | 0.01429 | 0.151 | 0.701 |
| | Percent hard food | 1 | 0.0774 | 0.07741 | 0.818 | 0.374 |
| | Dietary diversity | 1 | 0.0295 | 0.02951 | 0.312 | 0.581 |
| | Mean size | 1 | 0.0027 | 0.00271 | 0.029 | 0.867 |
| | SVL | 1 | 0.0342 | 0.03419 | 0.361 | 0.553 |
| | Percent plant: Mean size | 1 | 0.0177 | 0.01775 | 0.188 | 0.668 |
| | Percent hard food: Mean size | 1 | 0.002 | 0.00199 | 0.021 | 0.886 |
| | Dietary diversity: Mean size | 1 | 0.1102 | 0.11018 | 1.165 | 0.29 |
| | Residuals | 26 | 2.4593 | 0.09459 | | |

Table 7. Results from a reduced dataset of 35 toothrows: ANCOVAs investigating the tooth complexity, diet and tooth size, with SVL added as a covariate to control for ontogenetic stage. Asterisk indicates significant p-value.

| | Variable | Df | Sum Sq | Mean Sq | F-value | P-value |
|---|---|---|---|---|---|---|
| Average cusp number~ | Percent plant | 1 | 0.4811 | 0.4811 | 10.007 | 0.00394* |
| | Percent hard food | 1 | 0.0771 | 0.0771 | 1.604 | 0.21662 |
| | Dietary diversity | 1 | 0.0105 | 0.0105 | 0.218 | 0.64476 |

| | | | | | | |
|---|---|---|---|---|---|---|
| | Mean size | 1 | 0.0119 | 0.0119 | 0.247 | 0.62303 |
| | Tooth positions | 1 | 0.0025 | 0.0025 | 0.052 | 0.82132 |
| | Percent plant: Mean size | 1 | 0.0054 | 0.0054 | 0.113 | 0.73908 |
| | Percent hard food: Mean size | 1 | 0.0209 | 0.0209 | 0.434 | 0.51575 |
| | Dietary diversity: Mean size | 1 | 0.0811 | 0.0811 | 1.687 | 0.20539 |
| | Residuals | 26 | 1.2499 | 0.0481 | | |
| Mean tooth complexity~ | Percent plant | 1 | 2.642 | 2.6418 | 7.904 | 0.00925* |
| | Percent hard food | 1 | 0.042 | 0.0422 | 0.126 | 0.72522 |
| | Dietary diversity | 1 | 0.012 | 0.0119 | 0.036 | 0.85184 |
| | Mean size | 1 | 0.024 | 0.0239 | 0.071 | 0.79144 |
| | Tooth positions | 1 | 0.07 | 0.0701 | 0.21 | 0.65067 |
| | Percent plant: Mean size | 1 | 1.115 | 1.1154 | 3.337 | 0.07923 |
| | Percent hard food: Mean size | 1 | 0.704 | 0.7042 | 2.107 | 0.1586 |
| | Dietary diversity: Mean size | 1 | 0.043 | 0.0425 | 0.127 | 0.72425 |
| | Residuals | 26 | 8.69 | 0.3342 | | |
| Max tooth complexity~ | Percent plant | 1 | 1.381 | 1.381 | 1.406 | 0.2465 |
| | Percent hard food | 1 | 0.471 | 0.471 | 0.479 | 0.4949 |
| | Dietary diversity | 1 | 1.248 | 1.248 | 1.27 | 0.2701 |
| | Max size | 1 | 4.778 | 4.778 | 4.863 | 0.0365* |
| | Tooth positions | 1 | 1.919 | 1.919 | 1.954 | 0.174 |
| | Percent plant: Max size | 1 | 0.005 | 0.005 | 0.005 | 0.9443 |
| | Percent hard food: Max size | 1 | 1.269 | 1.269 | 1.292 | 0.2661 |
| | Dietary diversity: Max size | 1 | 0.002 | 0.002 | 0.002 | 0.9661 |
| | Residuals | 26 | 25.545 | 0.982 | | |
| Toothrow standard deviation~ | Percent plant | 1 | 0.0143 | 0.01429 | 0.162 | 0.69 |
| | Percent hard food | 1 | 0.0774 | 0.07741 | 0.879 | 0.357 |
| | Dietary diversity | 1 | 0.0295 | 0.02951 | 0.335 | 0.568 |
| | Mean size | 1 | 0.0027 | 0.00271 | 0.031 | 0.862 |
| | Tooth positions | 1 | 0.1767 | 0.17674 | 2.008 | 0.168 |
| | Percent plant: Mean size | 1 | 0.0763 | 0.07626 | 0.866 | 0.361 |
| | Percent hard food: Mean size | 1 | 0.0007 | 0.00072 | 0.008 | 0.928 |
| | Dietary diversity: Mean size | 1 | 0.0808 | 0.08077 | 0.917 | 0.347 |
| | Residuals | 26 | 2.2889 | 0.08803 | | |

Supplementary Table 8. Results from a reduced dataset of 35 toothrows: ANCOVAs investigating the tooth complexity, diet and tooth size, with tooth positions added as a covariate to control for ontogenetic stage. Asterisk indicates significant p-value.

| Extent of herbivory | Cohen’s d | Hedges’ g | Effect size classification |
|---|---|---|---|
| Minimal: Low | -0.1075115 | -0.1032111 | Very small |
| Low: Medium | 0.9787825 | 0.9290139 | Large |
| Medium: High | -0.02087654 | -0.01954399 | Very small |
| Minimal: High | -0.9943604 | -0.9470099 | Large |

Table 9. Results from a reduced dataset of 35 toothrows: tests investigating effect size between different categories of herbivore. Effect size classification is according to Sawilowsky, (2009), based on Hedge’s g value.

| Amount of hard food | Cohen’s d | Hedges’ g | Effect size classification |
|---|---|---|---|
| Very soft: Soft | 2.380604 | 2.214516 | Large |
| Soft: Medium | 1.195607 | 1.119292 | Large |

| | | | |
|---|---|---|---|
| Medium: High | 0.9929212 | 0.9424337 | Large |
| High: Very high | 0.6552513 | 0.6219334 | Medium |
| Very soft: Very high | 2.096968 | 1.863972 | Large |

Supplementary Table 10. Results from a reduced dataset of 35 toothrows: tests investigating effect size between different categories of hardness of diet. Effect size classification is according to Sawilowsky, (2009), based on Hedge's g value.
